# KSR2 promotes self-renewal and clonogenicity of small-cell lung carcinoma

**DOI:** 10.1101/2022.02.11.480157

**Authors:** Dianna H. Huisman, Deepan Chatterjee, Robert A. Svoboda, Heidi M. Vieira, Abbie S. Ireland, Sydney Skupa, James W. Askew, Danielle E. Frodyma, Luc Girard, Kurt W. Fisher, Michael S. Kareta, John D. Minna, Trudy G. Oliver, Robert E. Lewis

**Affiliations:** Eppley Institute, Fred & Pamela Buffett Cancer Center, University of Nebraska Medical Center, Omaha, NE 68198, USA; Department of Pathology, Microbiology, and Immunology, University of Nebraska Medical Center, Omaha, NE 68198, USA; Deparment of Molecular Cancer Biology, Duke University, Durham, NC, USA; Hamon Center for Therapeutic Oncology Research, Department of Pharmacology, Simmons Cancer Center, University of Texas Southwestern Medical Center, Dallas, Texas; Cancer Biology and Immunotherapies Group, Sanford Research, Sioux Falls, South Dakota; Department of Internal Medicine, University of Texas Southwestern Medical Center, Dallas, Texas; Deparment of Pharmacology and Cancer Biology, Duke University, Durham, NC, USA

## Abstract

Small-cell lung carcinoma (SCLC) tumors are heterogeneous, with a subpopulation of cells primed for tumor initiation. Here, we show that Kinase Suppressor of Ras 2 (KSR2) promotes the self-renewal and clonogenicity of SCLC cells. KSR2 is a molecular scaffold that promotes Raf/MEK/ERK signaling. KSR2 is preferentially expressed in the ASCL1 subtype of SCLC (SCLC-A) tumors and is expressed in pulmonary neuroendocrine cells, one of the identified cells of origin for SCLC-A tumors. The expression of KSR2 in SCLC and pulmonary neuroendocrine cells (PNECs) was previously unrecognized and serves as a novel model for understanding the role of KSR2-dependent signaling in normal and malignant tissues. Disruption of KSR2 in SCLC-A cell lines inhibits the colony forming ability of tumor propagating cells (TPCs) *in vitro* and their tumor initiating capacity *in vivo.* The effect of KSR2 depletion on self-renewal and clonogenicity is dependent on the interaction of KSR2 with ERK. These data indicate that the expression of KSR2 is an essential driver of SCLC-A tumor propagating cell function, and therefore may play a role in SCLC tumor initiation. These findings shed light on a novel effector promoting initiation of ASCL1-subtype SCLC tumors, and a potential subtype-specific therapeutic target.

**Implications:** Manipulation of the molecular scaffold KSR2 in ASCL1-subtype small-cell lung cancer cells reveals its contribution to self-renewal, clonogenicity, and tumor initiation.

## Introduction

Small-cell lung carcinoma (SCLC) affects current and former heavy smokers, accounting for 13% of all lung cancers^1^. There have been few improvements in SCLC detection, treatment, and survival in almost 40 years, leading to its classification as a recalcitrant cancer in 2012^1^. The five-year relative survival rates for SCLC patients with localized, regional, and distant disease are 30%, 18%, and 3%, respectively (American Cancer Society, 2024). Currently, SCLC patients are treated with first line therapy (cisplatin or carboplatin combined with etoposide chemotherapy, plus anti-PDL1 antibody, atezolizumab), second (topotecan), and third line (PD1 antagonist, nivolumab) therapies, and for some patients thoracic radiation therapy^2^. Although SCLC tumors are responsive to therapy initially, residual disease quickly develops resistance leading to the low five-year survival^3^. Substantial efforts have been made to characterize SCLC tumors and identify targets that may be selectively toxic to tumor cells while preserving normal lung tissue^4–8^. Rigorous and innovative basic science using state-of-the-art genetically-engineered mouse (GEM) models, and an extensive set of cell lines have led to key discoveries regarding the cells-of-origin and the common recurring mutations that underlie SCLC^7, 9–18^. These discoveries have yielded comprehensive genomic profiles and a durable classification of SCLC subtypes based on the differential expression of four key transcription factors, ASCL1 (SCLC-A), NEUROD1 (SCLC-N), POU2F3 (SCLC-P) and YAP1 (SCLC-Y)^7, 13, 17, 18^. Recently it has been suggested that the YAP1 subtype may not be a distinct subtype and can be reclassified as undifferentiated SMARCA4-deficient malignancies^19^. A fourth subtype of SCLC characterized by expression of ATOH1 has been reported^20^, with ATOH1 also being implicated in neuroendocrine differentiation of lung adenocarcinomas^21^.

Lineage tracing and single-cell RNA sequencing (scRNA-seq) in genetically modified mice showed that a rare pulmonary neuroendocrine cell (PNEC) subpopulation, NE^stem^ cells, actively responds to lung injury of the epithelia by expanding, migrating, and undergoing Notch-dependent transit amplification to regenerate the damaged epithelium^22^. This effort additionally identified NE^stem^ as a cell-of-origin for SCLC following Trp53, Rb1 and Notch mutations causing constitutive activation of stem cell renewal and altered deprogramming^22^. SCLC tumors in some GEM models have a small population of tumor propagating cells (TPCs) essential to the initiation, long-term propagation, and metastatic capacity of the tumor, while the bulk non-TPC population is highly proliferative but incapable of establishing tumors *in vivo*^23^. TPCs have been implicated in the initiation and growth of SCLC as well as therapy resistance^24–30^. TPCs are also implicated in epithelial-to-mesenchymal transition (EMT) and metastasis^30^. Their slower cycling and self-renewing ability enhances DNA repair, rendering these cells resistant to DNA damage-dependent chemo and radiation therapy^26, 27, 30,31^. Thus, SCLC TPCs offer a unique population within which to search for new targets, which in combination with current standard-of-care therapies may yield a durable and effective strategy for therapy.

Kinase Suppressor of Ras proteins KSR1 and KSR2 are molecular scaffolds for Raf/MEK/ERK signaling^32, 33^. While KSR1 is widely expressed, KSR2 expression is restricted to the brain, pituitary, adrenal glands, and neuroendocrine tissues^34, 35^. KSR2 is abundantly expressed in the brain, and its disruption reduces body temperature, promotes cold intolerance, impairs glucose homeostasis, elevates fasting insulin and free fatty acid levels in the blood, and causes obesity^33^. ChIP-seq analysis revealed *KSR2* as a transcriptional target in human ASCL1-subtype SCLC (SCLC-A) cell lines^36, 37^. KSR2 was identified as one of 24 druggable and overexpressed target genes of ASCL1 identified by ChIP-seq, providing a rationale for studying the role of KSR2 in SCLC-A^38^. Our work shows that KSR2 is expressed in PNECs and SCLC-A tumors and cell lines. Expression of KSR2 promotes the self-renewal and clonogenicity of SCLC-A TPCs and their tumor initiating capacity *in vitro* and *in vivo*. While KSR2 disruption is rescued by an intact KSR2 transgene, the inability of an ERK-binding deficient KSR2 construct to restore clonogenicity and tumor initiation implicates its scaffolding function in TPC formation. This result defines a novel mechanism of tumor initiation in SCLC-A and a potential therapeutic vulnerability.

## Materials and Methods

### Cell culture

Murine small-cell lung carcinoma cell lines KP1 and KP3 were a gift from J. Sage (Stanford University). Human small-cell lung carcinoma cell lines H209 and H1963 were a gift of John Minna (UT Southwestern). The cells were cultured in RPMI 1640 medium containing 10% fetal bovine serum (FBS) and grown at 37°C with ambient O_2_ and 5 % CO_2_. Cells were passaged a minimum of twice after thawing prior to use in experiments. Serum starved cells were grown in above conditions, with serum removed for 16 hours prior to cell lysis. Serum shocked cells were grown normally followed by boosting FBS concentration to 50% for 15 minutes prior to cell lysis. Ionomycin (ThermoFisher Scientific I24222) treatment was given at a 1 µM dose for 15 minutes prior to cell lysis. All cells were routinely tested for mycoplasma and confirmed negative, with the most recent test performed 1/19/25. No further authentication of cell lines was performed by the authors.

### Generation of KSR2 shRNA knockdown and CRISPR/Cas9 knockout cell lines

Individual SMARTvector human inducible lentiviral shRNAs suitable for targeting murine and human KSR2 expressed in piSMART hEF1a/TurboGFP vector were stably transfected into KP1, KP3, and H209 SCLC cell lines with polyethylenimine hydrochloride molecular weight 40,000 (PEI, Polysciences Inc., cat. 24765). Cells were selected for expression of the shRNAs using 0.25 µg/mL of puromycin. 48 hours after doxycycline induction, cells were selected again by flow cytometry sorting for GFP+ cells. Knockdown of KSR2 was confirmed by western blot. KSR2 cDNA (MSCV KSR2 IRES YFP) and ERK binding deficient KSR2 cDNA (MSCV KSR2 FIF570 IRES YFP) was made resistant to binding of hairpin sh5 by introducing three point mutations in the binding region. Point mutations were introduced using the QuikChange Lightning Site Directed Mutagenesis Kit (Agilent #210518) according to the manufacturer’s protocol. Sequencing of the construct confirmed the correct point mutations were made with no additional mutations. MSCV KSR2 IRES YFP resistant to binding hairpin sh5 (KSR2r) and MSCV KSR2 FIF570 IRES YFP resistant to binding hairpin sh5 (FIF570r) were transfected into HEK-293T cells using trans-lentiviral packing system (ThermoFisher Scientific). The virus was collected 48 hours post transfection and used to infect KP1 sh5 cells with 8 µg/mL Polybrene for 72 hours. KP1 sh5 cells expressing the KSR2r or FIF570r construct were selected for using flow cytometry sorting YFP+ cells. Presence of the KSR2r or FIF570r expression after doxycycline induced downregulation on endogenous KSR2 was confirmed via western blotting. To generate knockout cells, sgRNA sequences targeting KSR2 or non-targeting control (NTC) were inserted into lentiCRISPR v2 (Addgene #52961). The constructs were PEI transfected into HEK293T cells along with psPAX2 lentiviral packaging construct (Addgene #12259) and pMD2.G envelope construct (Addgene #12259). Lentivirus-containing media was harvested at 72-h and used to infect the H209 and H2107 cells with 8 µg/mL Polybrene. Cells were selected for expression of the sgRNAs using 2 ug/mL of puromycin, and western blot was used to confirm KSR2 knockout.

### Cell lysis, western blot analysis, and immunoprecipitation

Whole cell lysate was extracted in radioimmunoprecipitation assay (RIPA) buffer containing 50 mM Tris-HCl, 1% NP-40, 0.5 % Na deoxycholate, 0.1 % Na dodecyl sulfate, 150 mM NaCl, 2 mM EDTA, 2 mM EGTA, and protease and phosphatase inhibitors aprotinin (0.5 U/mL), leupeptin (20 mM), and NA_3_VO_4_ (0.5 mM). The estimation of protein concentration was done using BCA protein assay (Promega #PI-23222, PI-23224). Samples were diluted using 1 X sample buffer (4 X stock, LI-COR #928–40004) with 100 mM dithiothreitol (DTT) (10 X stock, 1 mM, Sigma #D9779-5G). The protein was separated using 8–12% SDS-PAGE and transferred to nitrocellulose membrane. The membrane was blocked with Odyssey TBS blocking buffer (LICOR-Biosciences #927–50003) for 45 min at room temperature, then incubated with primary antibodies (*Key Resources Table, Supplementary Table 1*) at least overnight at 4°C. IRDye 800CW and 680RD secondary antibodies (LI-COR Biosciences # 926–32211, # 926–68072) were diluted 1:10,000 in 0.1% TBS-Tween and imaged on the Odyssey Classic Scanner (LI-COR Biosciences). Immunoprecipitation was performed with precoated anti-FLAG beads (Sigma #M8823) overnight at 4°C. Samples were washed with three times with TBS, then eluted off beads by heating to 95°C in 1X sample buffer and 100 mM DTT and western blot was performed as above.

### Analysis of KSR2 transcript expression in normal tissues and mouse tumors

GTEx portal was used to display the relative expression of human *KSR2* mRNA (TPM) in brain-cortex and lung tissue. In a murine model, neuroendocrine specific reporter, *Chga*-GFP was used to identify PNECs. GFP+ neuroendocrine cells from 3 mouse lungs were isolated by flow cytometry. qPCR was performed to measure mRNA expression of *Ksr2*, *Cgrp*, *Chga*, *Syp*, and *Spc* in GFP+ neuroendocrine cells and GFP-lung epithelial cells. RPR2, RPM and RPMA mouse tumor data is derived from Ireland *et al.*, Cancer Cell, 2020^39^ and Olsen *et al*., Genes Dev, 2021^40^ and deposited in NCI GEO: GSE149180 and GSE155692.

### SCLC sequencing analysis

RNA sequencing data from human primary tumor samples^13^ was analyzed for *KSR2* expression based on high and low *ASCL1* expression or high and low *MYC* expression. RNA sequencing data of human SCLC cell lines (DepMap Portal, Broad Institute MIT) was segregated by SCLC subtype and analyzed for *KSR2* mRNA levels. RNA samples from SCLC (n = 69) and NSCLC-NE (n = 14) cell lines were analyzed by paired-end RNA sequencing as previously described (McMillan *et al*., 2018, Cell 173, 864–878; dbGaP study accession: phs001823.v1.p1). Briefly, reads were aligned to the human reference genome GRCh38 using STAR-2.7 (https://github.com/alexdobin/STAR [github.com]) and FPKM values were generated with cufflinks-2.2.1 (http://cole-trapnell-lab.github.io/cufflinks/ [cole-trapnell-lab.github.io]). All data were then pooled, normalized to Transcripts Per Millions (TPM), and log-transformed. RNAseq from the SCLC primary tumors (George *et al*., 2015, Nature 524, 47-53) were included in the analysis.

### Fluorescence Activated Cell Sorting

For live cell staining, SCLC cell lines were incubated 20 minutes on ice in PBS with DAPI (3 µM), PE-CD24 (1:400) PE-Cy7-CD44 (1:300) and APC-EpCAM (1:100). Cells were resuspended in PBS and flow cytometry of SCLC cell lines was performed using a 100 µm nozzle on a BD FACSAria II using FACSDiva software. Debris were excluded by gating on forward scatter area versus side scatter area. Doublets were excluded by gating on forward scatter area versus side scatter height. Viable cells were identified by exclusion of DAPI stained cells. CD24^high^CD44^low^ cells were included by sequential gating followed by EpCAM^high^ TPCs. Compensation was performed using single stain and fluorescence minus one (FMO) controls. Positive gates were set based on the negative unstained sample. Data were analyzed using FlowJo software. For fixed cell staining, SCLC cell lines were fixed in 4% formaldehyde for 15 minutes at room temperature then washed in 1X PBS. Cells were permeabilized using 100% methanol at -20°C for 10 minutes, then blocked for 1 hour in 1X PBS / 5% normal goat serum / 0.3% Triton™ X-100. Cells were probed with Phospho-p44/42 MAPK antibody (Cell Signaling #9101S) 1:500 in 1X PBS / 1% BSA / 0.3% Triton X-100 overnight at 4°C. After washing in 1X PBS cells were incubated with secondary antibody (Jackson ImmunoResearch Laboratories #111-545-003) for 2 hours at room temperature in the dark. Cells were rinsed and staining of CD24, CD44, and EPCAM antibodies, and flow cytometry gating was performed similarly to live cell staining procedure described above.

### Colony Formation Assay

For colony formation assays, SCLC cells were dissociated by gentle pipetting. Live TPCs were sorted using a 100 μm nozzle on a BD FACSAria II. TPCs were sorted individually into 96 well plates filled with regular media (200 μl/well) containing DMSO or doxycycline (DOX) (1 µg/mL). 50 µL fresh media with or without DOX was added to the wells every 10 days. Three weeks later, colony numbers were assessed using CellTiter-Glo 2.0 reagent (Promega #G9242) and luminescence was measured (POLARstar Optima plate reader) according to the manufacturer’s protocol. CellTiter-Glo® readings greater than 300 Relative Luminescence Units (RLUs) in colony forming assays were considered colonies.

### Growth Curve

Cells were seeded at low density (500 cells/well) in 100 µl in replicate 96-well plates (Fisher #237105), with each condition plated in a single column of the 96 well plate. Each replicate plate was reserved to be read on a single day. Wells were fed with complete media or doxycycline containing media and assessed every other day for a period of 6 days using CellTiter-Glo® 2.0 reagent (Promega #G9242). Data were analyzed by non-linear regression using GraphPad Software, Inc.

### Extreme Limiting Dilution Analysis

#### In vivo

The viable cell number was assessed by replicate cell counts on a hemocytometer using Trypan Blue exclusion. Viable cell number was used to derive a titration of cell numbers for implantation. Cells were diluted in 50 μl media (RPMI +10% FBS) and 50 μl Cultrex PathClear BME (Trevigen # 3632-005-02). Six eight-week-old NCG mice were injected subcutaneously into the shoulders and flanks. 3 replicates for each dilution were used. Mice were provided drinking water with 5% sucrose, or sucrose plus doxycycline (2 mg/kg). Injection sites were palpated biweekly to monitor for tumor growth and all mice were sacrificed when any one tumor reached 1 cm. Tumors that formed were scored as 1 and the absence of tumor formation was scored as 0 for the extreme limiting dilution analysis (ELDA). Tumors that formed were analyzed for expression of KSR2 by western blot and tumors in the doxycycline treated group which did not maintain knockdown were scored as 0. ELDA software was used to estimate TPC frequency for control and doxycycline treated groups.

#### In vitro

Cells were seeded in 96-well plates (Fisher #12556008) at decreasing cell concentrations (1000 cells/well – 1 cell/well) at half log intervals (1000, 300, 100, 30, 10, 3, 1), 12 wells per condition. Cells were cultured for 14 days, and wells with spheroids >100 µm were scored as spheroid positive. TPC frequency and significance between groups was calculated by ELDA software.

### Tissue Staining

At the conclusion of *in vivo* ELDA, tumors were excised and formalin fixed. Samples were then paraffin embedded and stained for H&E and Ki67 (Abcam, ab16667, 1:200) by the University of Nebraska Medical Center Tissue Sciences Facility. The University of Nebraska Medical Center Advanced Microscopy Facility performed whole slide scanning using the Zeiss Axioscan 7 Whole Slide Imaging System. Images were cropped for display using ZEN lite software on a Zeiss workstation made available by UNMC Advanced Microscopy Core Facility.

### Data Availability Statement

The data generated in this study are available upon request from the corresponding author. RPR2, RPM and RPMA mouse tumor data is derived from Ireland *et al.*, Cancer Cell, 2020^39^ and Olsen *et al*., Genes Dev, 2021^40^ and deposited in NCI GEO: GSE149180 and GSE155692. RNAseq from the SCLC and NSCLC tumors and cell lines was derived from George *et al*., 2015, Nature 524, 47-53.

## Results

### Pulmonary neuroendocrine cells and ASCL1-subtype SCLC tumors express KSR2

ASCL1-subtype (SCLC-A) tumors can arise from PNECs^22, 36, 38, 41^. PNECs are heterogeneous, including a small subpopulation termed NE^stem^ cells that respond to injury of the lung epithelia by expanding, migrating to the location of the damage, and undergoing Notch-dependent transit amplification to regenerate the damaged epithelium^22^. SCLC tumors may arise from PNECs following *Trp53*, *Rb1*, and *Notch* mutations causing constitutive activation of stem cell renewal and altered deprogramming^22^. Although *KSR2* mRNA is not detectable in normal epithelial lung tissue (Fig. 1A), it is present in PNECs (Fig. 1B). Analysis of human SCLC tumors showed that ASCL1-high and MYC-low tumors^13^ preferentially express *KSR2* mRNA (Fig. 1C-E). We examined expression of *Ksr2* in an established *Ascl1*-high *Rb1/p53/Rbl2* (RPR2) model as well as a MYC-driven SCLC model (*Rb1/p53/Myc*, RPM) associated with reduced *Ascl1* and the acquisition of other SCLC subtypes including *Neurod1* and *Yap1,* and a MYC-driven tumor model lacking *Ascl1* (*Rb1/p53/Myc*/*Ascl1*, RPMA) (Fig. 1F-G)^39, 40^ (NCI GEO: GSE149180 and GSE155692). *Ksr2* was highly enriched in RPR2 tumors and significantly depleted in RPM tumors when *Ascl1* was genetically depleted (RPMA), demonstrating a dependency of *Ksr2* expression on ASCL1 (Fig. 1G). Consistent with the mouse tumor data, analysis of *KSR2* mRNA by subtype reveals that *KSR2* is highly expressed in SCLC-A, but has varying expression in NEUROD1, POU2F3, and YAP1 subtypes (Fig. 1H). While *KSR2* mRNA is preferentially expressed in SCLC-A compared to other subtypes^7^, *KSR1* mRNA is detected across SCLC subtypes^13^ (Supplementary Table S2). SCLC-A and -N cells express KSR1, but only SCLC-A cells express KSR2 by western blot analysis (Fig. 1I, 1J).

**Fig. 1.**
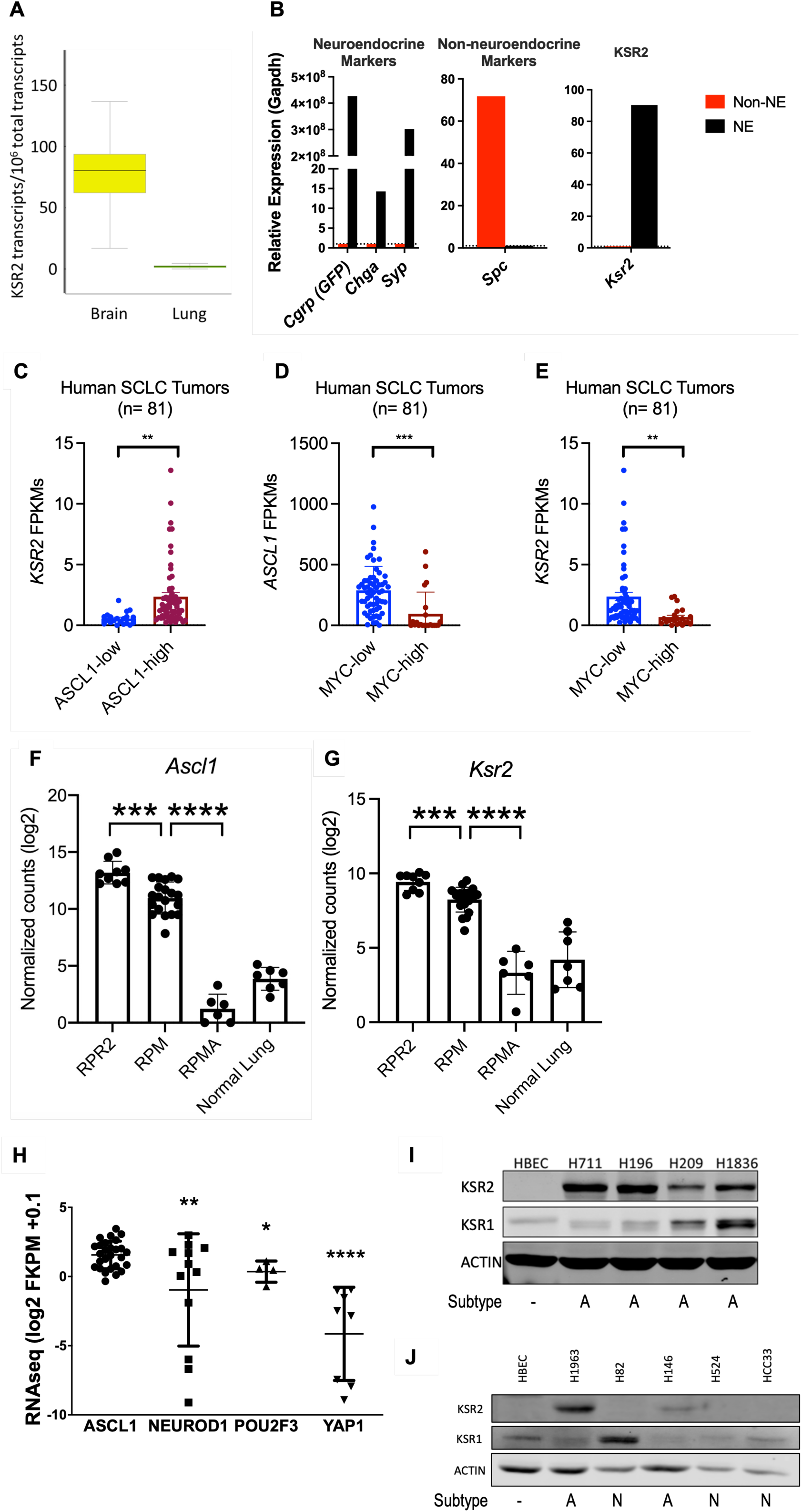
KSR2 expression across SCLC subtypes. A) *KSR2* mRNA in brain tissue and lung epithelial tissue (GTEx Portal). B) GFP_+_ cells isolated from the lungs of mice expressing GFP from the promoter for the neuroendocrine (NE)-specific *Cgrp.* mRNA expression of neuroendocrine markers (*Chga*, *Syp*) (*left*), lung epithelial marker surfactant protein C (*Spc*) (*middle*), and *Ksr2* (*right*). *KSR2* (C, E) and *ASCL1* (D) expression in human SCLC tumors based on *MYC* and *ASCL1* expression status. **, p<0.01 ***, p<0.001. *ASCL1* (F) and *Ksr2* (G) expression from bulk RNA-sequencing of indicated mouse tumors. H) *KSR2* RNA-seq (DepMap) in SCLC segregated by subtype. *p=0.03; **p<0.004; ****p<0.0001. I) KSR2 and KSR1 expression, with ACTIN loading control, in western blots of ASCL1-subtype SCLC cell lines and non-transformed human bronchial epithelial cells (HBEC). J) KSR2 and KSR1 expression, with ACTIN loading control, in western blots of ASCL1 (A) or NEUROD1 (N) SCLC lines and non-transformed human bronchial epithelial cells (HBEC).

### Depletion of Kinase Suppressor of Ras 2 reduces SCLC clonogenicity and self-renewal

SCLC-A tumor propagating cells (TPCs) are essential for tumor initiation and metastasis and are enriched by selection for high expression of CD24 and EpCAM, but low expression of CD44^23^. Transplantation assays show that TPCs are a minor but highly tumorigenic subpopulation of SCLC cells characterized by their colony forming activity *in vitro*, which measures their capacity for clonogenicity and self-renewal^23, 42^. Mouse SCLC cell lines KP1 and KP3 were derived from spontaneous mouse models of SCLC with knockout of Rb1, Trp53 (KP1) or Rb1, Trp53 and p130 (KP3), which replicate the ASCL1 subtype of SCLC tumors^7, 11^. KP1 cells can be stained for CD24^high^CD44^low^EpCAM^high^ to isolate TPCs (Fig. 2A, Supplementary Fig. S1A). KP1 and KP3 cells expressing one of three doxycycline (dox)-inducible shRNAs targeting *Ksr2* (sh5, sh6, sh7) were treated with or without doxycycline (Fig. 2B-C). Flow sorted CD24^high^CD44^low^EpCAM^high^ TPCs or EpCAM^low^ non-TPCs isolated by FACS were plated as single cells in 96-well plates to be analyzed for colony formation by CellTiter-Glo®. Colony formation, a measure of self-renewing capability of an individual cell^43–45^, was enriched in TPCs, while non-TPCs formed fewer and less robust colonies (Fig. 2D). Dox-induced targeting of *KSR2* in KP1 cells has little effect on the proportion of CD24^high^CD44^low^EpCAM^high^ TPCs detected by FACS (Supplementary Fig. S1B). Following downregulation of KSR2 expression, KP3 TPCs were analyzed for colony formation (Fig. 2E), and the percentage of wells with robust growing colonies was reduced by 56% and 38% with *Ksr2* RNAi by sh6 and sh7, respectively, which reflects their effectiveness at targeting *Ksr2* (Fig. 2C). In KP1 cells, colony formation was assessed with or without dox-induced RNAi by hairpin sh5 after TPC isolation, which inhibited colony formation by 86% (Fig. 2F). These data demonstrate that KSR2 is a significant contributor to the clonogenic and self-renewing properties of SCLC TPCs.

**Fig. 2.**
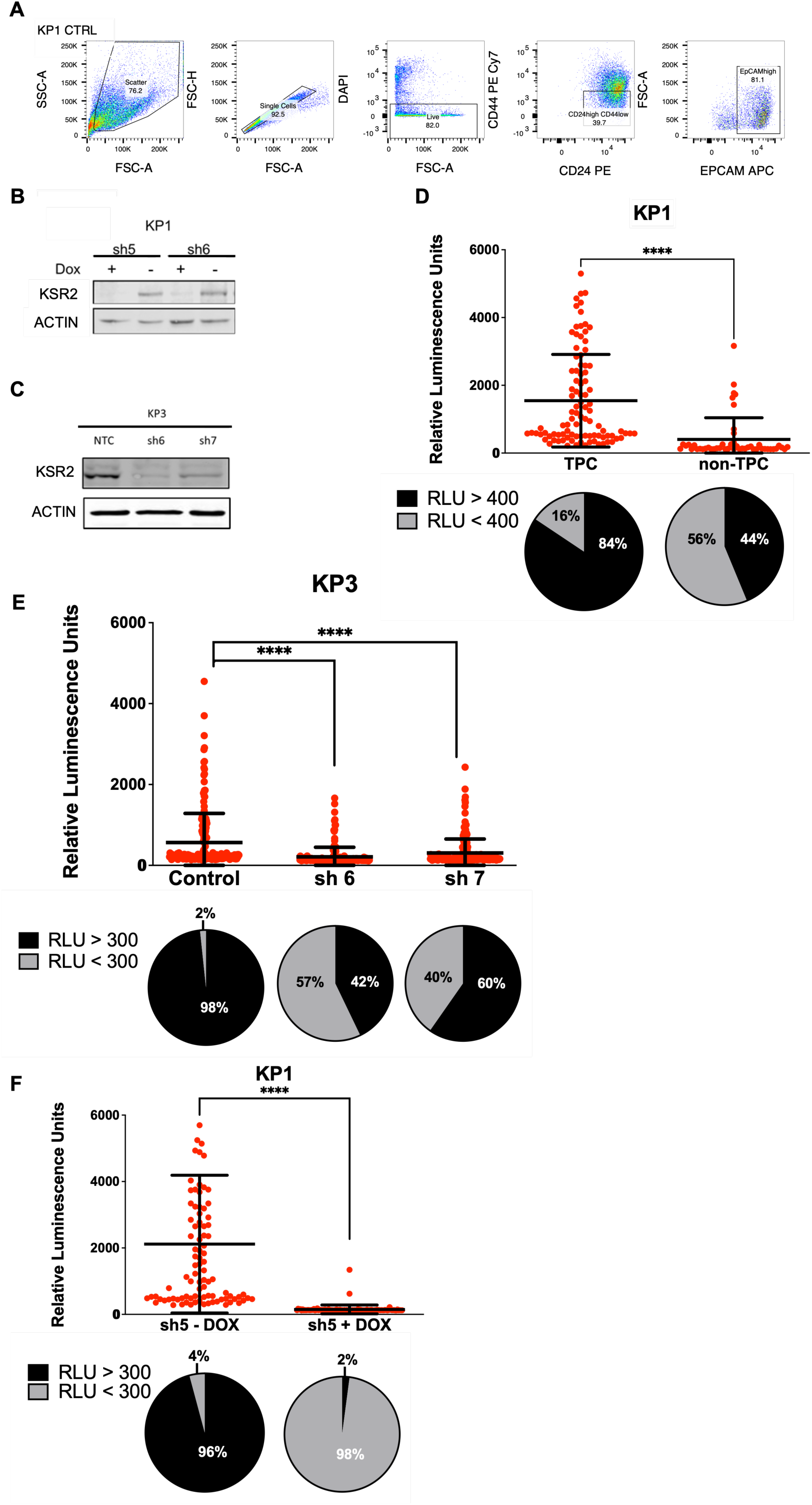
KSR2 disruption reduces tumor propagating cell clonogenicity in murine SCLC cells. A) FACS gating strategy of KP1 cells for Single, Live, CD24^high^ CD44^low^ and EPCAM^high^ TPCs. B-C) Dox-inducible RNAi of *Ksr2* (sh5, sh6, sh7) in KP1 and KP3 cells from an SCLC mouse model. NTC, non-targeting control. D) Clonogenicity of sorted CD24^high^ CD44^low^ and EPCAM^high^ TPCs versus EPCAM^low^ non-TPCs (above) and percentage of wells with robust colony formation (>400 RLU) (below). Dox-treated KP3 (E) and KP1 (F) SCLC TPCs with and without the indicated shRNAs targeting *Ksr2* (above) and percentage of colony forming wells (below). Colony viability was measured by CellTiter-Glo. ****, p<0.0001, n=3.

### Expression of a KSR2 transgene restores colony formation and increases TPC frequency

To confirm on-target effect of our inducible shRNA system for KSR2 knockdown, a construct containing wildtype *Ksr2* mutated to be resistant to binding hairpin sh5 (KSR2r) was expressed in KP1 sh5 cells (Fig. 3A). Rescue of KSR2 expression restored colony formation to wildtype levels in KP1 sh5 cells confirming the on-target effect of hairpin sh5 (Fig. 3B). *In vitro* ELDA was used to estimate the relative frequency of TPCs. KSR2 disruption reduced TPC frequency from 1/19 in control KP1 cells to 1/166 in KP1 cells with KSR2 KD (Fig. 3C, D). The expression of wildtype *Ksr2* construct KSR2r restored the TPC frequency to 1/18 (Fig. 3C, D). These data demonstrate the on-target action of the inducible *Ksr2* shRNA system to regulate TPC abundance.

**Fig. 3.**
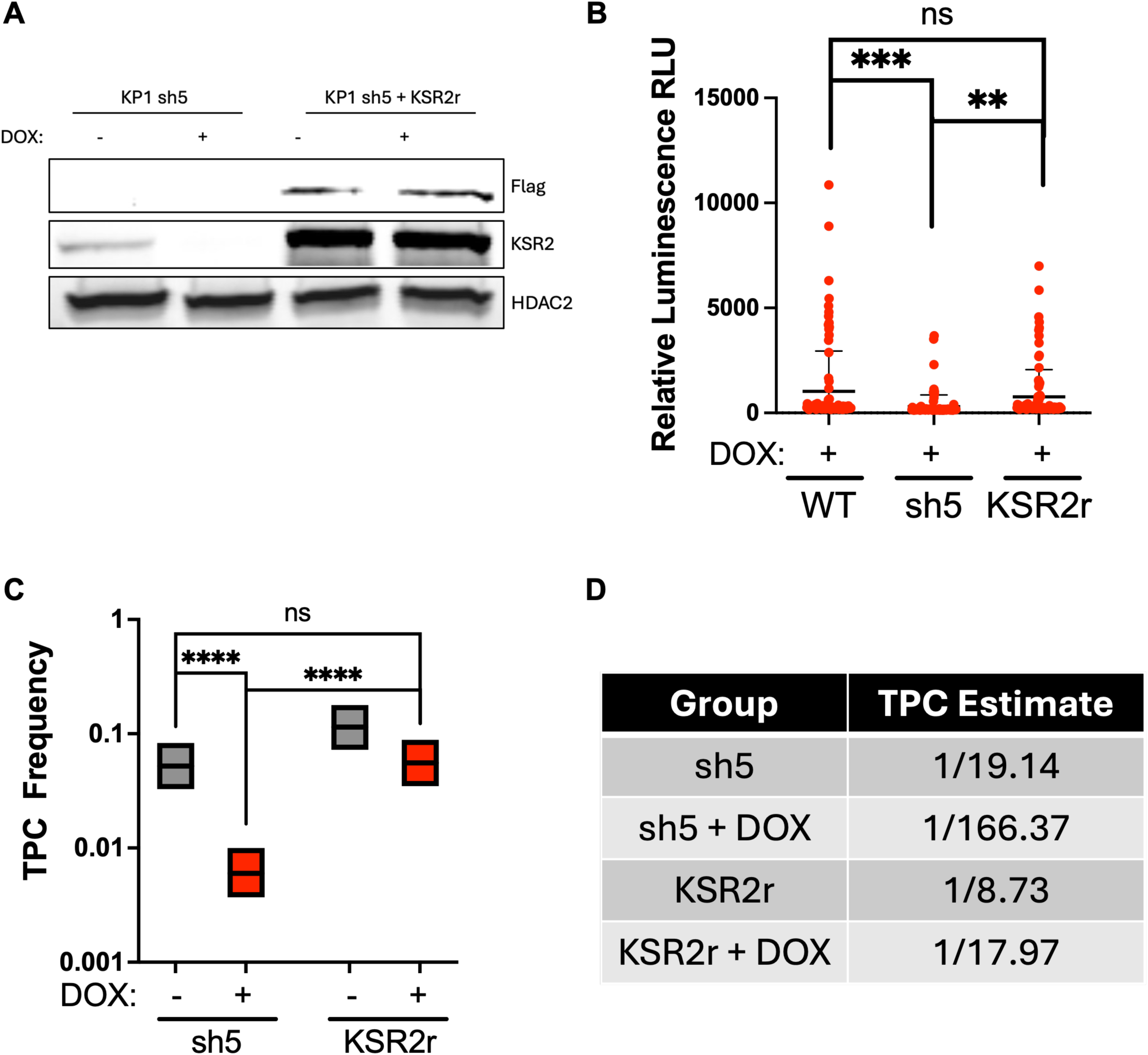
SCLC TPC frequency and clonogenicity is rescued by a KSR2 transgene. A) A *Ksr2* cDNA resistant to the sh5 shRNA (KSR2r) was introduced into KP1 cells expressing the Dox-inducible sh5 shRNA. B) Clonogenicity of KP1 SCLC TPC colonies within wildtype (WT), dox-induced RNAi of *Ksr2* (sh5) and dox-induced RNAi of *Ksr2* + expression of shRNA-resistant *Ksr2* (KSR2r). Colony viability was measured by CellTiter-Glo. ****, p<0.0001. C-D) *In vitro* ELDA of KP1 sh5 or KSR2r KP1 cells with dox-induced RNAi of KSR2 was used to determine the estimated frequency of TPCs. ****, p<0.0001.

### KSR2 disruption inhibits the tumor initiating capacity of human SCLC cell lines *in vitro*

*In vitro* ELDA was used to test the effect of doxycycline (dox)-induced *KSR2* disruption on human SCLC cell line H209 (Fig. 4A, C). The TPC frequency of control H209 cells was reduced 12-fold from 1/4 in control cells to 1/48 in H209 sh5 cells (Fig. 4C). KSR2 was also targeted using a CRISPR/Cas9 in conjunction with a *KSR2* sgRNA (sg2, sg3), or non-targeting control (NTC) in H209 and H2107 cells (Fig. 4B, E). The TPC frequency of H209 cells was reduced 4- to 7-fold from 1/44 in NTC to 1/187-1/302 in KSR2 KO cells (Fig. 4D). The TPC frequency of H2107 cells was reduced 4- to 5-fold from 1/37 in NTC to 1/149-1/174 in KSR2 KO cells (Fig. 4F) These data indicate that KSR2 supports tumor initiating capacity in human SCLC tumor propagating cells.

**Fig. 4.**
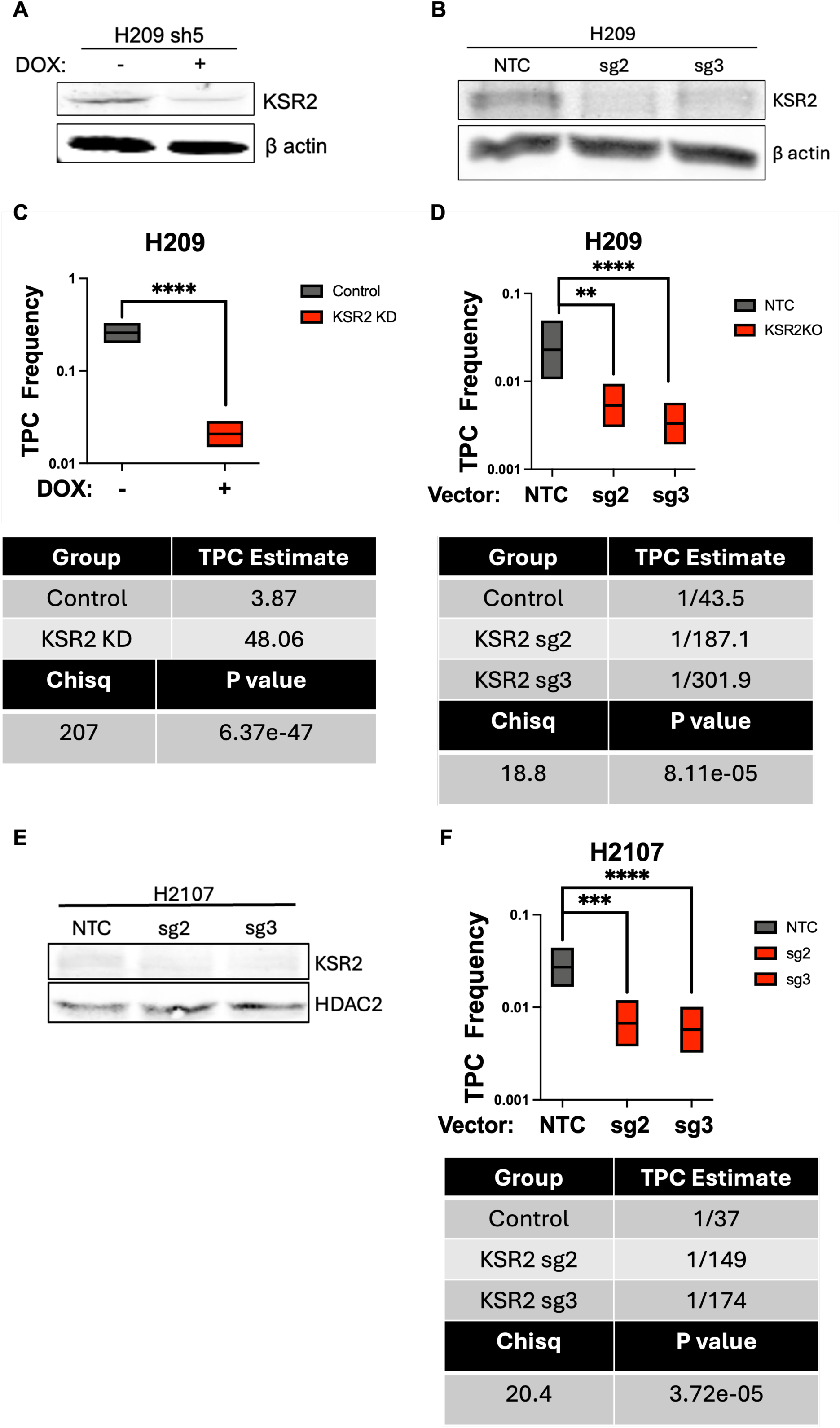
SCLC TPC frequency is reduced by KSR2 knockdown or knockout in human SCLC cells. A) Western blot of H209 cells with dox-induced RNAi of *KSR2* (sh5) or following CRISPR/Cas9 expression with non-targeting (NTC1) control gRNA or sgRNA2 or 3 (KSR2KO) targeting *KSR2* (B). C-D) *In vitro* ELDA was used to determine the proportion of TPCs in H209 cells with dox-induced RNAi of *KSR2* (sh5) or CRISPR/Cas9 targeting of *KSR2* (sg2, sg3). E) Western blot of H2107 cells following CRISPR/Cas9 expression with non-targeting (NTC1) control gRNA or sgRNA2 or 3 (KSR2KO) targeting *KSR2*. F) *In vitro* ELDA was used to determine the proportion of TPCs in H2107 cells with CRISPR/Cas9 targeting of *KSR2* (sg2, sg3). ****, p<0.0001. ***, p<0.001. **, p<0.01.

### KSR2 disruption inhibits the tumor initiating capacity of murine SCLC cells *in vivo*

Extreme limiting dilution analysis (ELDA) is a statistical method optimized for estimating the stem cell frequency from limiting dilution analysis^46^. Stem cell frequency within the bulk tumor cell population is determined from the frequency of tumor-positive and tumor-negative injections arising from varying doses of xenografted cells. Dox-inducible shRNA targeted KP1 cells were injected with successive dilutions into NOD-*Prkd^cem26Cd52^Il2rg^em26Cd22^*/NjuCrl (NCG) mice, and the mice were provided drinking water with sucrose, or sucrose plus doxycycline (2 mg/kg). ELDA was performed by scoring each tumor that arose as “1”. The absence of tumor formation was scored as “0”. KSR2 disruption reduced frequency of TPCs 10-fold, from 1/255 control KP1 cells to 1/2530 KP1 cells with KSR2 knockdown (KD, hairpin sh5) (Fig. 5A, C, Supplementary Table S3). ELDA was performed *in vivo* with H209 cells targeted by CRISPR/Cas9 for *KSR2* (sg2), with KSR2 knockout (KO) reducing TPC frequency 6-fold from 1/265 to 1/1602 (Fig. 5B, D, Supplementary Table S4). Isolated H209 control (NTC) and KSR2KO (sg2) tumors from the ELDA show robust Ki67 staining with minimal induction of PARP cleavage or Caspase 3 cleavage (Supplementary Fig. S2C, E). This is consistent with *in vitro* growth curves, in which KP1 cells with KSR2 KD and H209 cells with KSR2 knockout grow at the same rate as control cells (Supplementary Fig. S2A, B). These data indicate that KSR2 is required to illicit the full tumor initiation potential of SCLC tumor propagating cells *in vivo*. Analysis of subtype defining transcription factors in the isolated tumors revealed no strong effect, with a single KSR2 knockout tumor showing a mild downregulation of ASCL1 and coincident upregulation of NEUROD1 with no effect on POU2F3 expression (Supplementary Fig. S2D).

**Fig. 5.**
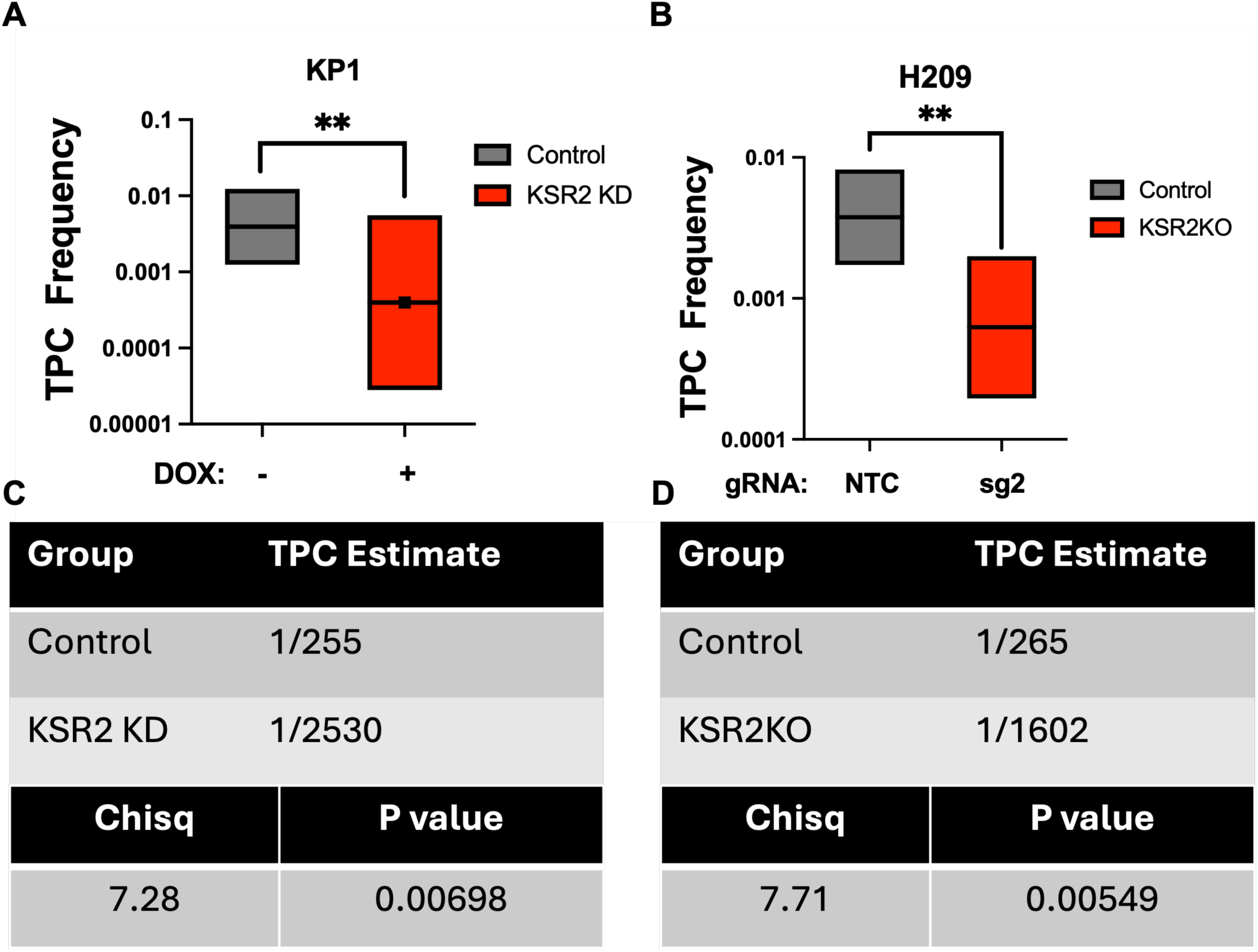
KSR2 disruption reduces frequency of tumor propagating cells *in vivo*. Raw data and quantification of *in vivo* extreme limiting dilution analysis of KP1 cells without and with Dox-induced RNAi of *Ksr2* (sh5) (A, C) **, p=0.00698 and H209 control (NTC) and KSR2 KO (sg2) cells (B, D) **, p=0.00549.

### KSR2-ERK interaction is necessary for SCLC clonogenicity

KSR2 knockdown reduces activation of ERK (Fig. 6A). Similarly, in serum starved and serum stimulated conditions KSR2 knockdown cells show reduced ERK activation (Fig. 6A). Cells expressing KSR2r rescue ERK activation in each condition (Fig. 6B). To assess the importance of the interaction between KSR2 and ERK, a DEF domain mutant (570 FIFP/AAAP 573)^32^ KSR2 construct deficient in its ability to bind ERK (FIF570) was further mutated to be resistant to binding hairpin sh5 (FIF570r) and expressed in KP1 sh5 cells. Cells expressing FIF570r are unable to rescue ERK activation when endogenous KSR2 is depleted (Fig. 6C). Disruption of KSR2/ERK interaction prevents ERK activation in response to serum stimulus. Immunoprecipitation of the FLAG epitope tag on this construct and the full length KSR2 demonstrated reduced phosphor-ERK associated with the ERK-binding mutant construct (FIF570r) (Fig. 6D).

**Fig. 6.**
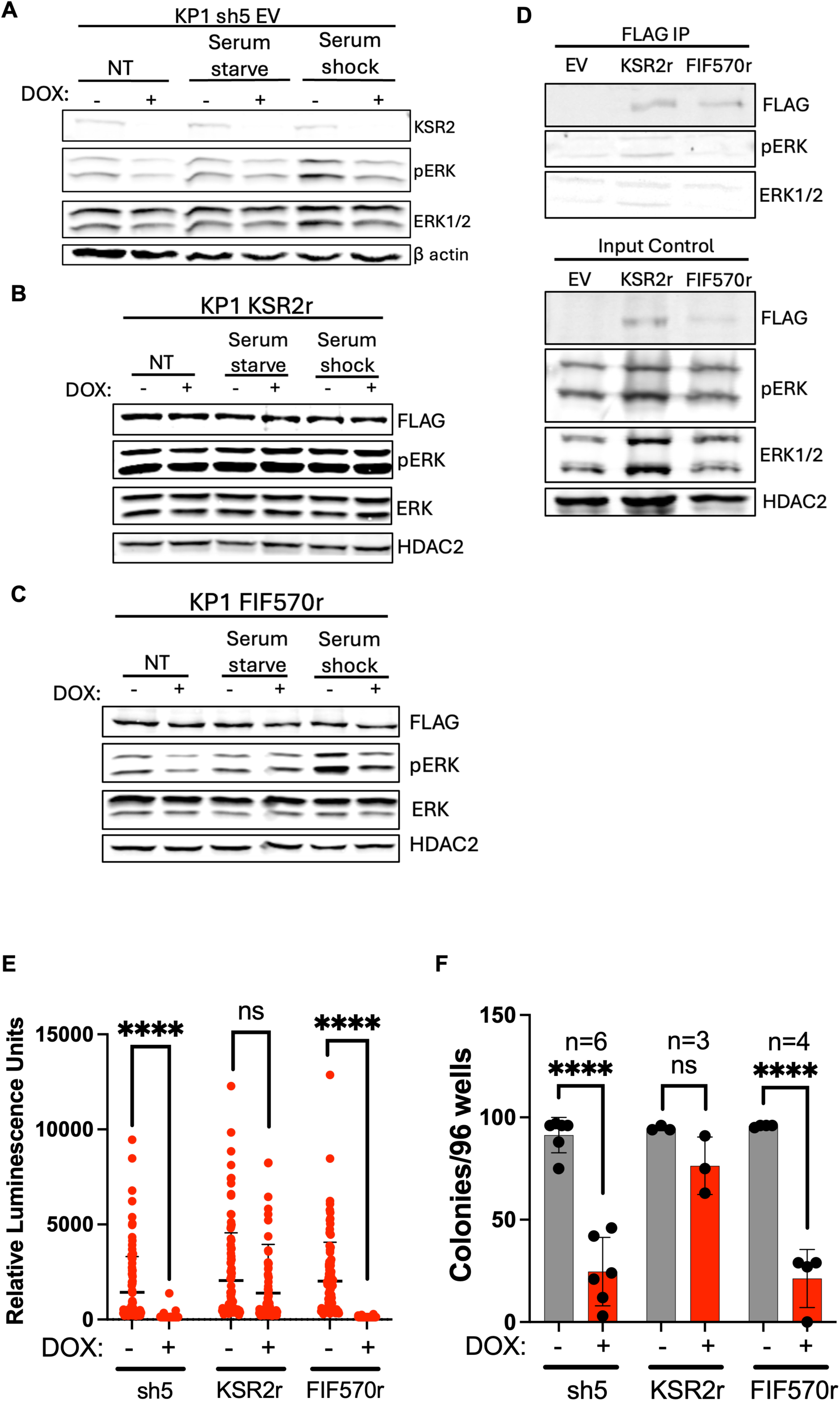
Disruption of KSR2-ERK interaction reduces clonogenicity of murine SCLC TPCs. Western blot showing phosphorylation of ERK in KP1 sh5 (A), FLAG epitope-tagged KSR2r (B), and FLAG epitope-tagged FIF570r (C) cells with and without dox-induced KSR2 depletion with no treatment (NT), serum starvation, or serum shock. D) Immunoprecipitation of FLAG epitope-tagged wildtype KSR2 (KSR2r) and ERK-binding mutant KSR2 (FIF570r), or empty vector control (EV). E) Clonogenicity of KP1 SCLC TPC colonies with dox-induced RNAi of *Ksr2* (sh5) and dox-induced RNAi of *Ksr2* + expression of hairpin resistant *Ksr2* (KSR2r) or hairpin resistant ERK-binding mutant *Ksr2* (FIF570r). F) Quantification of colonies greater than 300 RLU per 96/well plate, sample size is as indicated, ****, p<0.0001.

Colony formation was significantly reduced in the FIF570r cells after dox-induced targeting of endogenous KSR2, suggesting that KSR2 interaction with ERK is necessary for the clonogenic capacity of SCLC TPCs (Fig. 6E). Disruption of KSR2 significantly reduced the number of robust colonies formed by TPCs (Fig. 6F). KSR2r rescued colony formation (Fig. 6F), while FIF570r was unable to restore colony formation to KP1 TPCs (Fig. 6F). To better understand the importance of the interaction between KSR2 and ERK, and ERK activation for clonogenicity in TPCs, the percentage of phosphor-ERK high KP1 cells in bulk versus CD24^high^CD44^low^EpCAM^high^ TPC populations was analyzed and showed a statistically significant modest but modest 10% increase in the percentage of cells in the TPC population with high phosphor-ERK levels (Supplementary Fig. S3A-D). Due to the modest increase in ERK phosphorylation in CD24^high^CD44^low^EpCAM^high^ TPCs, ELDA was performed in KP1 cells in the presence of the MEK inhibitor Trametinib to determine if ERK phosphorylation had an impact of the frequency of TPCs. MEK inhibition reduced ERK phosphorylation (Supplementary Fig. S3F) but had no significant impact of TPC frequency (Supplementary Fig. S3E). These data indicate that the interaction between KSR2 and ERK is an important contributor to colony formation by TPCs, while ERK phosphorylation is not necessary for maintenance of the TPC population.

## Discussion

Our data reveal an unexpected role of KSR2 in ASCL1 subtype SCLC tumor initiation. These data suggest that KSR2 increases the number of tumor initiating in SCLC-A. Despite its ability to function as a Raf/MEK/ERK scaffold, the ability of KSR2 to enhance tumor initiation appears dependent upon its interaction with ERK but not on ERK activity.

Kinase Suppressor of Ras proteins KSR1 and KSR2 have unique and overlapping functions. Both function as scaffolds for Raf/MEK/ERK signaling^32, 33^ promoting phosphorylation of ERK. KSR2 is expressed in the brain, pituitary gland, and adrenal gland, and some neuroendocrine cells and tissues^34, 35, 47^. Pulmonary neuroendocrine cells are a cell-of origin for SCLC-A likely explaining the expression of KSR2 in this subtype, ASCL1 is a potential transcriptional activator of *KSR2*^38^, which may explain why KSR2 is not widely expressed in other SCLC subtypes. Here we show that disruption of KSR2 in SCLC-A reduces TPC clonogenicity and self-renewal *in vitro*, and the number of tumor initiating cells *in vivo*. Mutation of the KSR2 DEF domain, which is required for interaction with ERK^32^, demonstrates that the clonogenic capacity of TPCs depends upon the interaction of KSR2 with ERK. Activating Ras mutations are rare in SCLC tumors, therefore Raf/MEK/ERK signaling is likely activated by extracellular stimuli, suggesting that a molecular scaffold, such as KSR2, is required to amplify this signaling pathway^48^. The role of ERK signaling in SCLC is incompletely understood, but is implicated in cell proliferation, differentiation, survival, and drug resistance^49^.

Treatment with an ERK inhibitor was unable to induce apoptosis in human SCLC-A cell lines H209 and H69^50^. Our data contrast with observations that SCLC-A cell line proliferation was significantly reduced *in vitro* and in *in vivo* tumor xenografts by ARHGEF19 disruption and downstream reduced Raf/MEK/ERK signaling^51^. This conflict is interesting and deserves further investigation. Activation of Raf/MEK/ERK by endoplasmic reticulum (ER) stress has also been reported to promote SCLC cell survival^52^. Raf/MEK/ERK signaling may play an essential role in promoting metastasis of SCLC tumors^48^. CXCL12 induces ERK activation in SCLC cells, which correlates with increased invasion through extracellular matrix^53^. Increased Raf/MEK/ERK signaling induced by expression of an activated Raf construct has been reported to cause growth arrest and reduce expression of neuroendocrine markers in SCLC^54, 55^.

Our data reveal that KSR2 is important for self-renewal and clonogenicity of the SCLC-A TPC population and disruption of the ERK binding domain is sufficient to reduce their clonogenic capacity. We expected this effect to be due to the reduction in ERK activation and found that the percentage of cells with high ERK activation was modestly but significantly higher in the TPC population in comparison with the bulk cell population. However, it is difficult to attribute biological significance to this modest 10% increase in the number of TPCs with elevated phosphor-ERK. MEK inhibitor Trametinib reduces ERK phosphorylation without affecting the frequency of TPCs suggesting that the slight elevation of ERK phosphorylation is not necessary for clonogenic capacity. These observations demonstrate the complexity of ERK signaling in SCLC tumors, which may also be affected by the intensity, frequency, localization, or duration of ERK activation, and suggests that disruption of KSR2 may be a more effective therapeutic target than targeting Raf/MEK/ERK signaling directly. We may also consider that although ERK activation is not directly affecting the function of TPCs, that ERK may be acting as an allosteric regulator of KSR2 or that mutation of the DEF domain may disrupt KSR2 interaction with novel effectors. Future work testing these possibilities will be needed to detail the mechanisms through which KSR2 supports TPC function.

The effect of KSR2 depletion on ERK activation is surprising as SCLC-A cells express both KSR2 and KSR1. We do not detect KSR1 compensation for KSR2 loss, suggesting that KSR2 is required for TPC function in SCLC-A cells. Our data suggest that either KSR2 is contributing to TPC function by promoting signaling that KSR1 cannot influence, or that KSR2 and KSR1 coordinate function to conduct efficient signaling. KSR1 and KSR2 are capable of heterodimerization with Raf, resulting in a conformational change of KSR proteins that allows phosphorylation of MEK^56–58^. The dimerization of KSR proteins with Raf orients the Raf protein so that its catalytic site is not in close proximity to the phosphorylation site on MEK to complete the phosphorylation, which necessitates MEK phosphorylation by Raf through a *trans* interaction^56^. KSR2-BRAF heterodimerization results in an increase of MEK phosphorylation via the KSR2-mediated relay of signal from BRAF to release the activation segment of MEK for phosphorylation^56^. KSR2 can also homodimerize via the same interface that interacts with BRAF, however this creates a different quaternary structure when interacting with MEK and it’s unknown how this may affect the availability of MEK for phosphorylation^56^. BRAF homodimers, or KSR1-BRAF heterodimers, should generate a quaternary structure sufficient for availability of MEK for phosphorylation by an additional BRAF protein^56^. Via the same interaction interface, it is possible that KSR1 and KSR2 heterodimerize. Our incomplete understanding of how KSR2 homodimers or KSR1-KSR2 heterodimers may impact MEK phosphorylation makes it difficult to predict the effect of KSR2 loss on MEK phosphorylation and downstream ERK activation. However, KSR2 depletion has a deleterious effect on ERK activation in SCLC-A cells. It is notable that we fail to detect KSR2 protein expression in the NEUROD1-subtype of SCLC (SCLC-N), while they retain expression of KSR1 (Fig. 1E, F). This observation suggests that SCLC-N cells are capable of compensating for loss of KSR2 with KSR1 expression potentially implicating KSR1 as an important target in SCLC-N, or that the transition to the NEUROD1-subtype results in upregulation of other compensatory signaling pathways that negate the dependency on KSR2. Though KSR2 protein is not detected in these cells, targeting KSR1 in SCLC-N reduces tumor initiating cell frequency and enhances response to cisplatin (bioRxiv 2024.02.23.581815).

ASCL1 and NEUROD1 drive distinct transcriptional profiles in SCLC^14, 59, 60^. SCLC-A cells can be converted to SCLC-N cells by elevation of MYC^60^. MYC amplification has been associated with poor patient outcome, therapy resistance, and tumor progression^61–63^. KSR2 expression seems to be lost during transition from SCLC-A to SCLC-N. The initiation of switching from SCLC-A subtype to SCLC-N subtype is incompletely understood, although recent studies implicate that epigenetic mechanisms are involved^64^. The expression of KSR2 in SCLC-A is thought to be driven by its defining transcription factor ASCL1, as *KSR2* is a transcriptional target of ASCL1^36, 37^. In tumors, we report rare instances with KSR2 disruption in which ASCL1 expression is downregulated with a coincidence increase in NEUROD1 expression. To date, there is no evidence of KSR2 affecting expression of ASCL1, however its role in facilitating ERK activation may provide rationale for further examination of the effect of KSR2 dependent Raf/MEK/ERK signaling on SCLC subtype and neuroendocrine differentiation^65, 66^. ASCL1 and KSR2 are also expressed in SCLC-A cell-of-origin, PNECs. PNECs respond to injury of the lung epithelia by expanding, migrating, and undergoing Notch-dependent transit amplification to regenerate the damaged epithelium^22^. Although KSR2 plays a role in SCLC TPC function, its role in normal PNECs is undetermined. KSR2 knockout mice have metabolic defects^33, 34^ but have no reports of abnormal lung pathology. Studies determining the effect of KSR2 manipulation on lung repair are required to determine if KSR2 plays a role in the function of PNECs in response to damage.

If KSR2 is dispensable for normal lung repair, it represents a potential therapeutic target for SCLC-A tumors. The discovery that cell surface proteins CD24, CD44, and EpCAM can be used to enrich and isolate SCLC-A TPCs significantly advanced investigation of potential therapeutic vulnerabilities within this population^22^. Cisplatin treatment does not enrich TPCs indicating that TPCs are not a reservoir for drug tolerant persister cells^22^, however TPCs are capable of repopulating tumors after treatment suggesting that they are not preferentially sensitive to the chemotherapeutic. Isolation of TPCs allows for interrogation of potential therapies that may be selectively detrimental to the TPC population. The distinct transcriptional profiles of the SCLC subtypes are therapeutically relevant, as there are a number of MYC-associated vulnerabilities in ASCL1 low tumors which differ from vulnerabilities identified in ASCL1 high tumors^67^. Subtype specific vulnerabilities have been reported for all four SCLC subtypes^68^ suggesting that further investigation into subtype specific therapeutic options, and their clinical relevance, is necessary. The preferential expression of KSR2 in SCLC-A, and the role it plays in TPC function provide rationale for further interrogating KSR2 as a SCLC-A specific therapeutic target.

JQ1, an inhibitor of bromodomain and extra-terminal (BET) proteins, reduced the frequency of TPCs *in vivo* and suppressed the ability of single TPCs to form colonies *in vitro*^22^. Our data demonstrate that KSR2 disruption reduces frequency of TPCs and reduces colony formation *in vitro* and tumor initiation *in vivo*. The effect of KSR2 on TPC clonogenicity is dependent on interaction with ERK but independent of ERK phosphorylation. These results suggest that targeting of KSR2 or its interaction with ERK may be TPC-specific therapeutic strategies and may create a more durable therapeutic response when used in conjunction with standard of care chemotherapies. The data provides a rationale for interrogating the effects of targeting KSR2 in genetically engineered mouse models of SCLC-A^12, 14, 69^. KSR2 disruption may work well in combination with cisplatin to target TPCs, reducing tumor burden and recurrence.

## Acknowledgements

The authors thank the UNMC Flow Cytometry Research Facility for help isolating SCLC TPCs. The UNMC Flow Cytometry Research Facility is administrated through the Office of the Vice Chancellor for Research and supported by state funds from the Nebraska Research Initiative (NRI) and The Fred and Pamela Buffett Cancer Center’s National Cancer Institute Cancer Support Grant. Major instrumentation has been provided by the Office of the Vice Chancellor for Research, The University of Nebraska Foundation, the Nebraska Banker’s Fund, and by the NIH-NCRR Shared Instrument Program. We acknowledge use of the University of Nebraska Medical Center -UNMC Advanced Microscopy Core Facility, RRID:SCR_022467, P20 GM103427. We additionally thank James R. Talaska, B.S. of the UNMC AMCF for their assistance.

## Author Contributions

**D.H. Huisman:** Conceptualization, Data curation, Formal Analysis, Investigation, Methodology, Supervision, Validation, Visualization, Writing – original draft, Writing – review & editing; **D. Chatterjee:** Methodology, Validation, Writing – review & editing; **R.A. Svoboda:** Methodology; **H.M. Vieira:** Methodology; **A.S. Ireland:** Data curation, Formal Analysis, Methodology, Visualization; **S. Skupa:** Methodology; **J.W. Askew:** Methodology; **D.E. Frodyma:** Methodology; **L. Girard:** Data curation, Formal Analysis; **K.W. Fisher:** Methodology; **M.S. Kareta:** Data curation, Methodology; **J.D. Minna:** Funding acquisition, Resources, Writing – review & editing; **T.G. Oliver:** Funding acquisition, Resources, Visualization, Writing – review & editing; **R.E. Lewis:** Conceptualization, Data curation, Funding acquisition, Project administration, Resources, Supervision, Writing – original draft, Writing – review & editing

