## Supplementary Figure S1 for "KSR2 promotes self-renewal and clonogenicity of small-cell lung carcinoma"

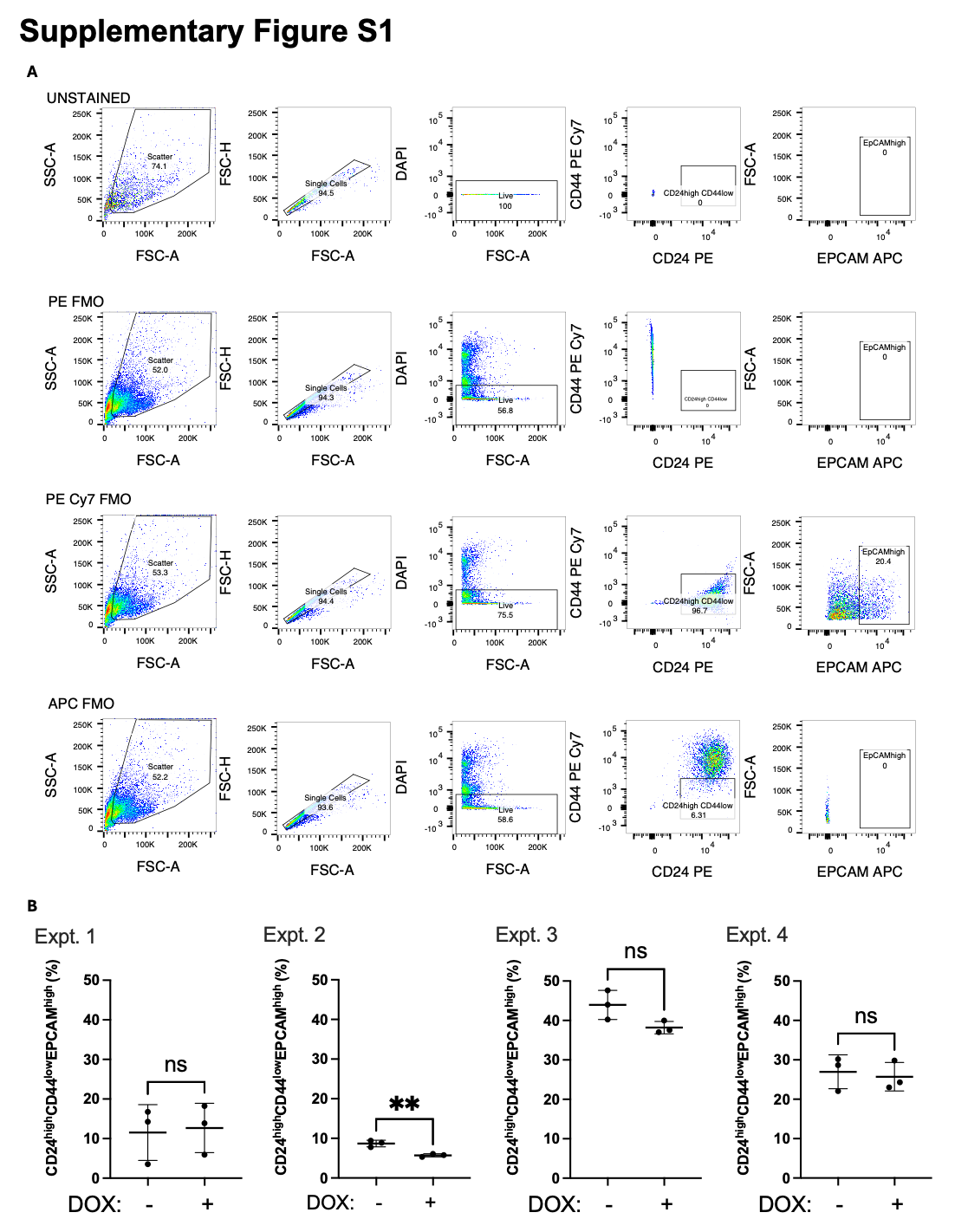

**Supplementary Figure S1. Sorting of CD24^high^CD44^low^EPCAM^high^ tumor propagating cells.** A) Flow cytometry control samples including unstained and fluorescence-minus-one (FMO) controls demonstrate gating scheme. B) Quantification of four biological replicate experiments measuring percentage of live KP1 cells that are CD24^high^CD44^low^EPCAM^high^ without (-) or with (+) 72 hour doxycycline treatment.
