## Supplementary Figure S2 for "KSR2 promotes self-renewal and clonogenicity of small-cell lung carcinoma"

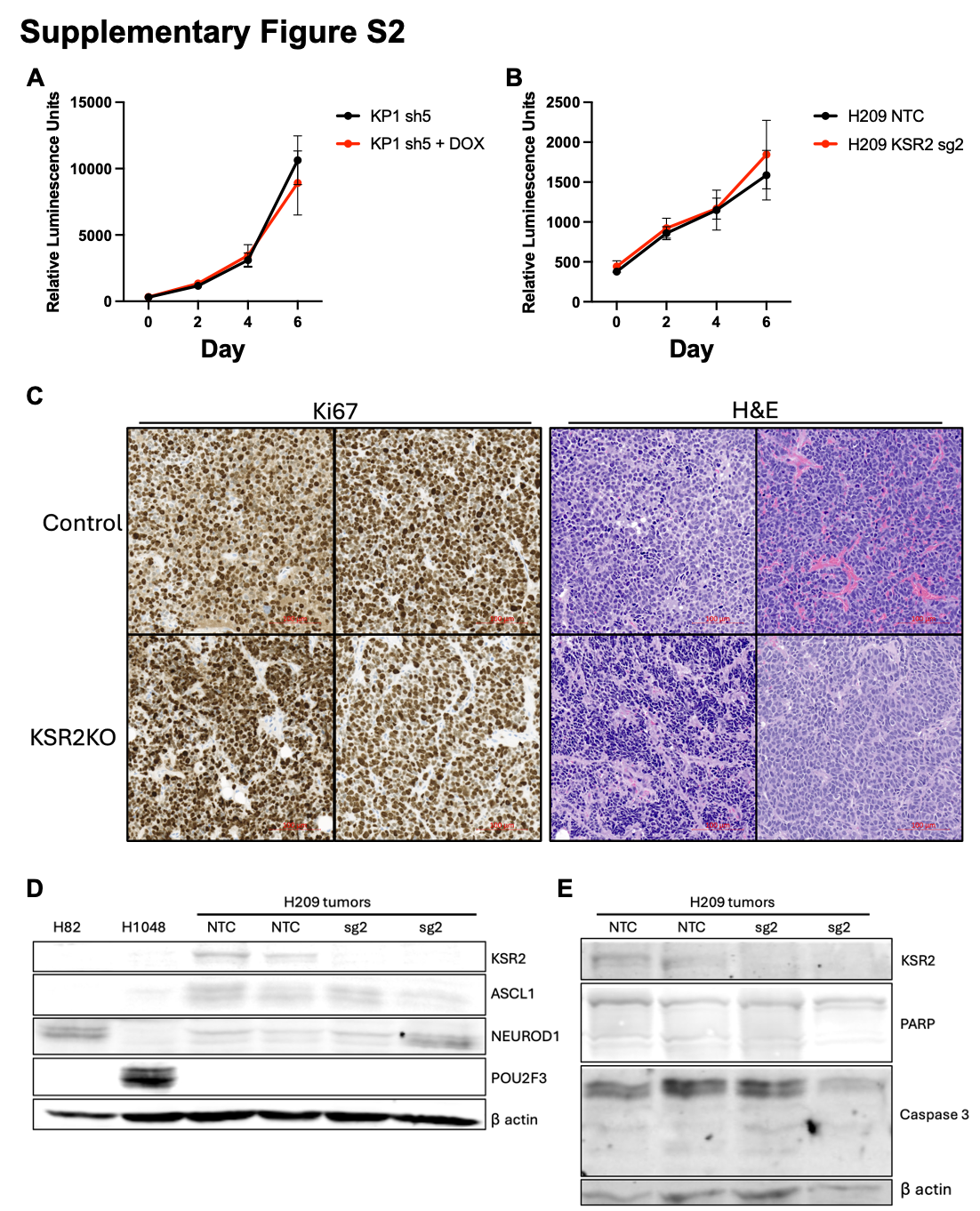

**Supplementary Figure S2. Proliferation, apoptosis, and subtype are not affected by KSR2 disruption in H209 tumor xenografts.** A) Growth curve of KP1 cells with or without Dox-induced targeting of *Ksr2* (sh5). B) Growth curve of H209 cells with CRISPR/Cas9 targeting of KSR2 (sg2) or non-targeting control (NTC). C) Ki67 and H&E staining in two replicate control and KSR2KO (sg2) samples of H209 tumor xenografts. D) Western blot showing SCLC subtype of H82, H1048, and H209 tumor xenografts with control (NTC) and KSR2KO (sg2). E) Western blot showing apoptosis markers in H209 tumor xenografts with control (NTC) and KSR2KO (sg2).
