## Supplementary Figure S3 for "KSR2 promotes self-renewal and clonogenicity of small-cell lung carcinoma"

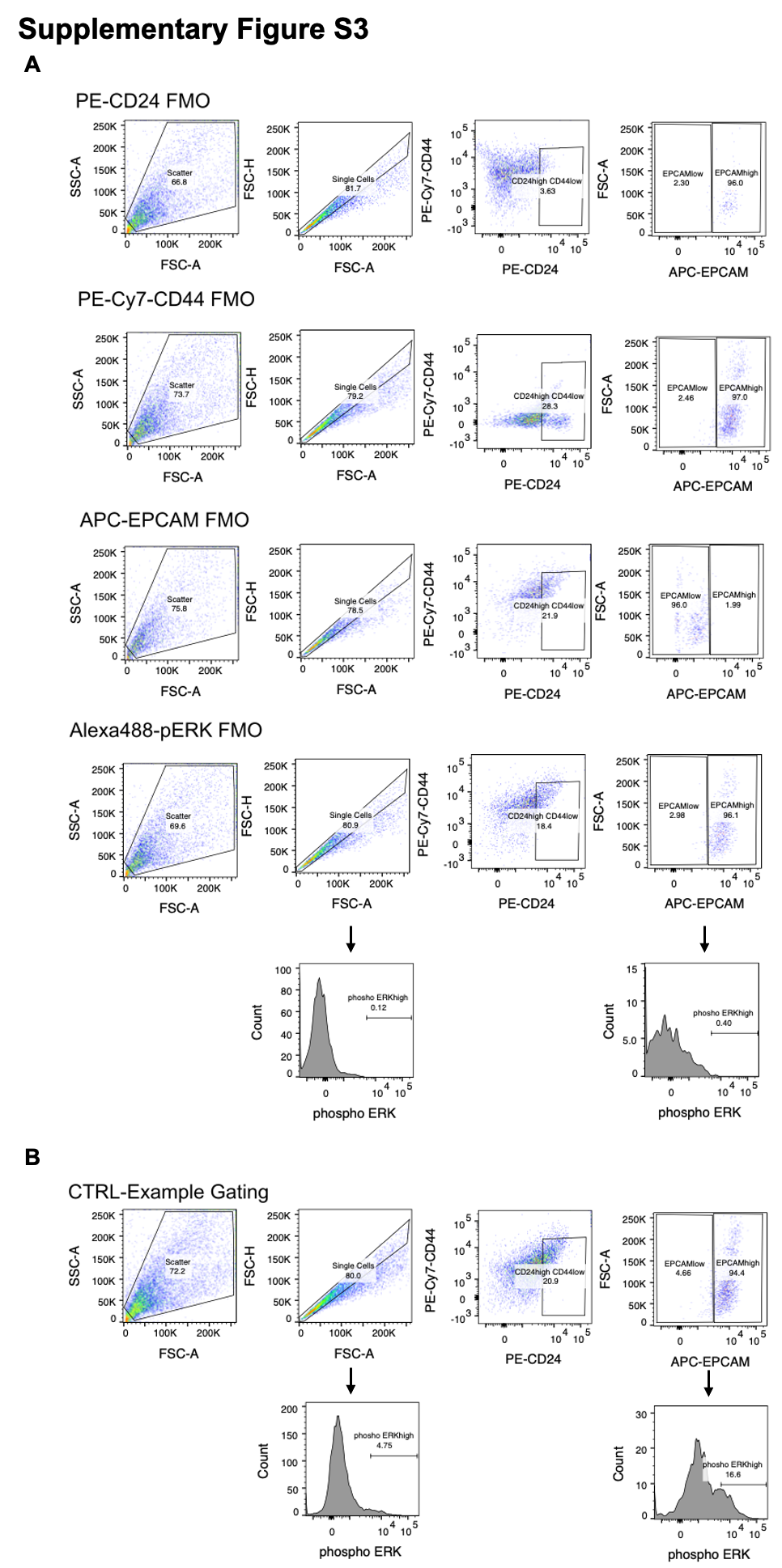

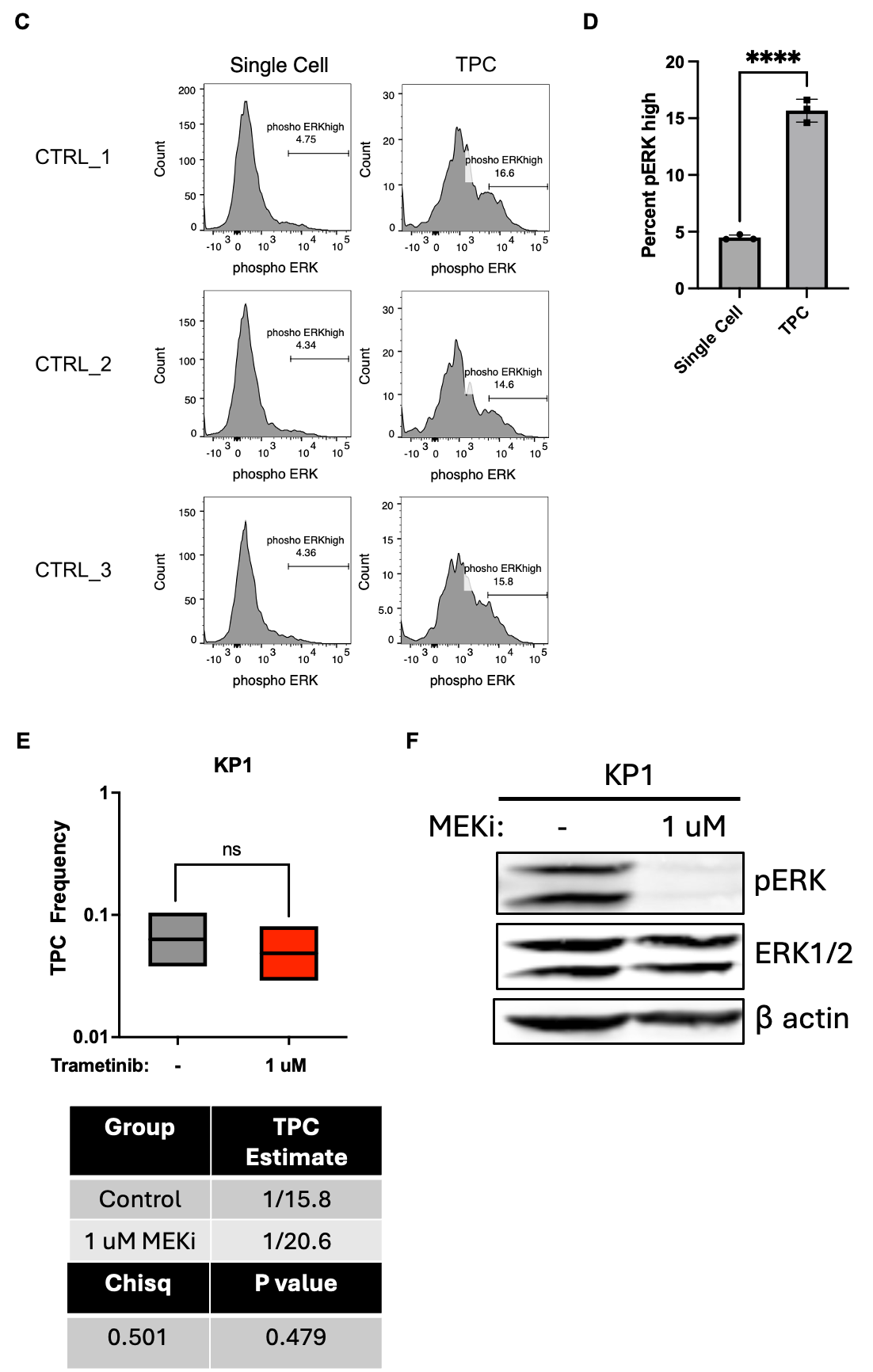

**Supplementary Figure S3. Inhibition of ERK phosphorylation does not affect TPC function.** A) FMO controls demonstrating gating scheme to identify AlexaFluor488-phosphoERK levels in Single or CD24^high^CD44^low^EPCAM^high^ KP1 cells. B) Example gating scheme comparing pERK levels in bulk (Single Cells) population versus TPC (CD24^high^CD44^low^EPCAM^high^) population. C) Histogram depicting pERK expression in bulk (single cell) versus CD24^high^CD44^low^EPCAM^high^ (TPC) population in three replicate samples. D) Quantification of percent pERK high cells in Single Cell versus TPC population. E) *In vitro* ELDA was used to determine the proportion of TPCs in KP1 cells ± 1 µM MEK inhibitor trametinib treatment. F) Western blot showing reduced ERK activation in response to 1u

M MEK inhibitor trametinib treatment.
