## Supplementary Table S1 for "KSR2 promotes self-renewal and clonogenicity of small-cell lung carcinoma"

**Key Resources**

| **Reagent type (species) or resource** | **Designation** | **Source or reference** | **Identifiers** | **Additional information** |
| --- | --- | --- | --- | --- |
| Cell line (*Mus musculus*) | Small-cell lung carcinoma | Obtained from Julien Sage (Stanford University) | KP1 |  |
| Cell line (*Mus musculus*) | Small-cell lung carcinoma | Obtained from Julien Sage (Stanford University) | KP3 |  |
| Cell line (*Homo sapiens*) | Small-cell lung carcinoma | Obtained from John Minna (UT Southwestern) | NCI-H209  RRID: CVCL_1525 |  |
| Cell line (*Homo sapiens*) | Small-cell lung carcinoma | Obtained from John Minna (UT Southwestern) | NCI-H1963  RRID: CVCL_1510 |  |
| Cell line (*Homo sapiens*) | Small-cell lung carcinoma | Obtained from John Minna (UT Southwestern) | NCI-H711  RRID: CVCL_1580 |  |
| Cell line (*Homo sapiens*) | Small-cell lung carcinoma | Obtained from John Minna (UT Southwestern) | NCI-H196  RRID: CVCL_1509 |  |
| Cell line (*Homo sapiens*) | Small-cell lung carcinoma | Obtained from John Minna (UT Southwestern) | NCI-H1836  RRID: CVCL_1498 |  |
| Cell line (*Homo sapiens*) | Small-cell lung carcinoma | Obtained from John Minna (UT Southwestern) | NCI-H82  RRID: CVCL_1591 |  |
| Cell line (*Homo sapiens*) | Small-cell lung carcinoma | Obtained from John Minna (UT Southwestern) | NCI-H146  RRID: CVCL_1473 |  |
| Cell line (*Homo sapiens*) | Small-cell lung carcinoma | Obtained from John Minna (UT Southwestern) | NCI-H524  RRID: CVCL_1568 |  |
| Cell line (*Homo sapiens*) | Small-cell lung carcinoma | Obtained from John Minna (UT Southwestern) | HCC33  RRID: CVCL_2058 |  |
| Cell line (*Homo sapiens*) | Small-cell lung carcinoma | Obtained from John Minna (UT Southwestern) | HBEC |  |
| Transfected construct | piSMART hEF1a/TurboGFP-KSR2 shRNA 5 | Dharmacon | Cat# V3SH11252-225640245 | GGAGCAAATCCCACGAGTT |
| Transfected construct | piSMART hEF1a/TurboGFP-KSR2 shRNA 6 | Dharmacon | Cat# V3SH11252-227968197 | GCACATCAGTCAGACGCTC |
| Transfected construct | piSMART hEF1a/TurboGFP-KSR2 shRNA 7 | Dharmacon | Cat# V3SH11252-228320208 | CCATCAAGCACAGGTTTTC |
| Transfected construct | lentiCRISPR v2- KSR2 sgRNA3 | Addgene | Plasmid #52961  RRID: Addgene_52961 | KSR2_3: 5’ CCGAATCGTCGATGTGCGCA 3’ |
| Transfected construct | lentiCRISPR v2- KSR2 sgRNA2 | Addgene | Plasmid #52961  RRID: Addgene_52961 | KSR2_2: 5’ GCTTACCGTGGACGCCTACC 3’ |
| Transfected construct | lentiCRISPR v2- NTC | Addgene | Plasmid #52961  RRID: Addgene_52961 | NTC: 5’ CCATATCGGGGCGAGACATG 3’ |
| Recombinant DNA reagent | KSR2r | (adapted from Fernandez et al. 2012)^29^ | MSCV KSR2 IRES YFP | QuikChange Lightning Mutagenesis: Forward 5’ CGCTGCACAGGAGTAAGTCCCAGTAATTCCAGCTCGGG 3’  Reverse 5’CCCGAGCGTGAATTCATGGGACTTACTCCTGTGCAGCG 3’ |
| Recombinant DNA reagent | FIF570r | (adapted from Fernandez et al. 2012)^29^ | MSCV KSR2 FIF570 IRES YFP | QuikChange Lightning Mutagenesis: Forward 5’ CGCTGCACAGGAGTAAGTCCCAGTAATTCCAGCTCGGG 3’  Reverse 5’CCCGAGCGTGAATTCATGGGACTTACTCCTGTGCAGCG 3’ |
| Sequence based reagent | Ksr2 (PCR primer) | (Guo et al. 2017)^31^ |  | Forward primer 5′-TGGATGTCCGAAAGGAAGTC-3′  Reverse primer 5′-CTTCTCCACGGTCTCACACA-3′ |
| Sequence based reagent | Cgrp (PCR primer) | (Lim et al. 2017)^38^ |  |  |
| Sequence based reagent | Chga (PCR primer) | (Lim et al. 2017)^38^ |  |  |
| Sequence based reagent | Syp (PCR primer) | (Lim et al. 2017)^38^ |  |  |
| Sequence based reagent | Spc (PCR primer) | (Lim et al. 2017)^38^ |  |  |
| Antibody | Anti-B ACTIN, mouse monoclonal | Santa Cruz | Cat# 47778  RRID: AB_626632 | WB 1:1000 |
| Antibody | Anti-KSR2 (MO8), mouse monoclonal | Abnova | Cat# H00283455-M08  RRID: AB_509404 | WB 1:1000 |
| Antibody | Phospho-p44/42 MAPK (Erk1/2) (Thr202/Tyr204) (E10), mouse monoclonal | Cell Signaling | Cat# 9106  RRID: AB_331768 | WB 1:1000  FACS: 1:500 |
| Antibody | p44/42 MAPK (Erk1/2), rabbit polyclonal | Cell Signaling | Cat# 9102  RRID: AB_330744 | WB 1:1000 |
| Antibody | Anti-Flag (M2), mouse monoclonal | Sigma | Cat# F3165  RRID: AB_259529 | WB 1:1000 |
| Antibody | Anti-HDAC2 (Y461), rabbit monoclonal | Abcam | Cat# ab32117  RRID: AB_732777 | WB: 1:1200 |
| Antibody | Anti-PARP, rabbit polyclonal | Cell Signaling | Cat# 9542  RRID: AB_2160739 | WB 1:1000 |
| Antibody | Anti-Caspase-3, rabbit polyclonal | Cell Signaling | Cat# 9662  RRID: AB_331439 | WB 1:1000 |
| Antibody | APC anti-mouse CD326 (EpCAM), mouse monoclonal | Biolegend | Cat# 118213  RRID: AB_1134105 | FACS: 1:100 |
| Antibody | PE anti-mouse CD24, mouse monoclonal | Biolegend | Cat# 101807  RRID: AB_312840 | FACS: 1:400 |
| Antibody | PE Cyanine7 anti-mouse/human CD44, mouse monoclonal | Biolegend | Cat# 103029  RRID: AB_830786 | FACS: 1:300 |
| Antibody | Alexa Fluor® 488 AffiniPure™ Goat Anti-Rabbit IgG (H+L) | Jackson ImmunoResearch Laboratories Inc. | Cat# 111-545-003  RRID: AB_2338046 | FACS: 1:500 |
| Antibody | Recombinant Anti-Ki67 | Abcam | Cat# Ab16667 RRID: AB_302459 | IHC: 1:200 |
| Nuclear Stain | DAPI (4',6-Diamidino-2-Phenylindole, Dilactate) | Biolegend | Cat#  422801 | FACS: 3 µM |
| NCG mouse | NOD-*Prkdc^em26Cd52^Il2rg^em26Cd22^*/NjuCrl | Charles River | Strain: 572  RRID:IMSR_CRL:572 |  |

**Supplementary Table S1. Key resources and reagents used in this study.**
