## Supplementary Table S3 for "KSR2 promotes self-renewal and clonogenicity of small-cell lung carcinoma"

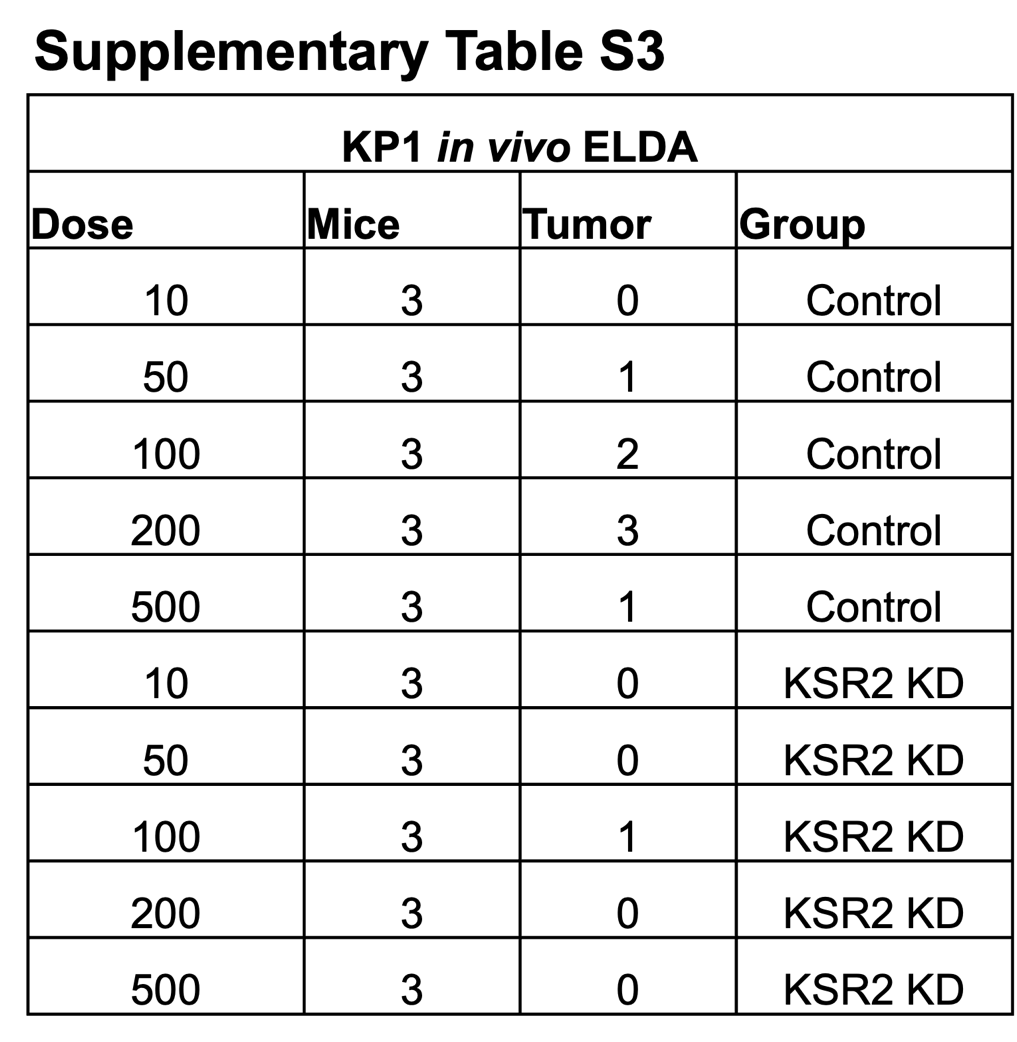

**Supplementary Table S3. Tumor formation in KP1 *in vivo* ELDA.** Number of tumors formed (Tumor) in three replicate mice across injections of different cell dilutions (Dose) of KP1 sh5 cells with (KSR2KD) or without (Control) administration of doxycycline by drinking water.
