## Supplementary Table S4 for "KSR2 promotes self-renewal and clonogenicity of small-cell lung carcinoma"

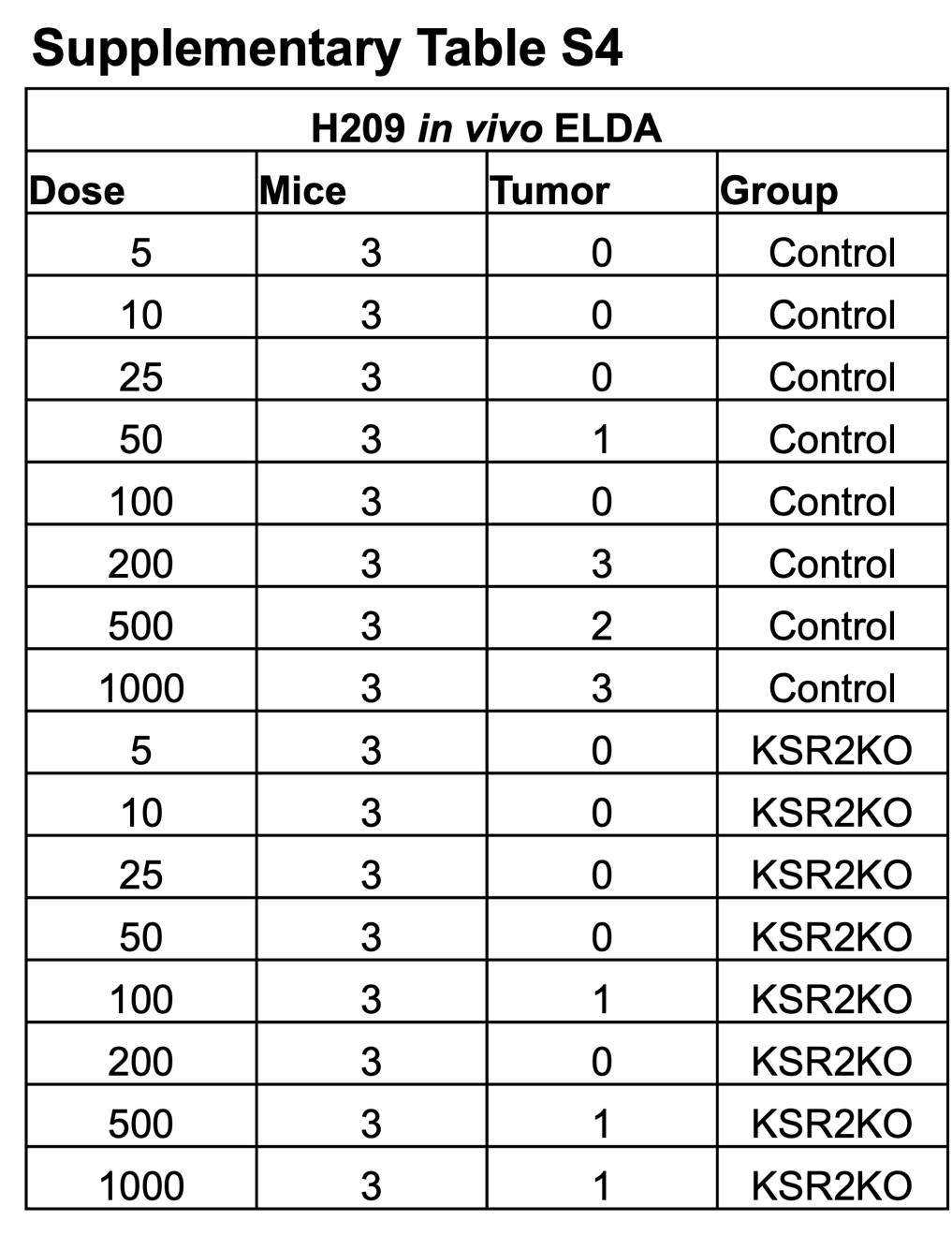

**Supplementary Table S4. Tumor formation in H209 *in vivo* ELDA.** Number of tumors formed (Tumor) in three replicate mice across injections of different cell dilutions (Dose) of H209 cells targeted by CRISPR/Cas9 with non-targeting control (Control) or *KSR2*-targeting sg2 (KSR2KO).
